# TIAM1 signaling drives prostatic budding and branching phenotypes and is a potential therapeutic target for BPH

**DOI:** 10.1101/2024.02.02.578055

**Authors:** Hamed Khedmatgozar, Sayanika Dutta, Michael Dominguez, Daniel Latour, Melanie Johnson, Mohamed Fokar, Irfan Warraich, Werner de Riese, Allan Haynes, Robert J. Matusik, Luis Brandi, Srinivas Nandana, Manisha Tripathi

## Abstract

Benign prostatic hyperplasia (BPH) is the most prevalent urologic disease in men aged over 50 years. However, the molecular mechanisms that drive BPH pathophysiology remain elusive. In this study, we integrated bioinformatic and experimental analyses of human BPH to identify TIAM1-RAC1 signaling pathway as a promising candidate for a molecular-based approach for BPH therapy. First, elevated TIAM1 expression in a BPH transcriptomic signature that was generated from the analysis of RNA-seq data from three independent BPH patient cohorts was validated at the protein level in a fourth patient cohort. Additional bioinformatic analyses of the BPH transcriptomic signature pointed to TIAM1-RAC1 pathway as the potential lead therapeutic pathway; and NSC23766 - a small molecule inhibitor of TIAM1 signaling - as a developmental lead compound for BPH therapy. Next, a proof-of-concept pharmacological approach of TIAM1-RAC1 inhibition in human prostatic cells using NSC23766 resulted in attenuated organoid budding and branching - a developmental program associated with prostatic nodule formation and BPH pathogenesis. Finally, shRNA-based genetic knock-down of TIAM1 in human prostatic cells led to a reduction in budding and branching phenotypes thereby phenocopying the effects of NSC23766. Together, our observations implicate elevated TIAM1 as a driver of budding and branching in BPH, and our studies pave the way for TIAM1-RAC1 based targeted approach for the treatment of the disease.

## Introduction

Benign prostatic hyperplasia (BPH) is a common pathological condition that affects more than half of men in their sixth decade, and >90% over 90 years of age. BPH causes significant symptoms and morbidity by obstructing urine outflow (also called bladder outlet obstruction) resulting in lower urinary tract symptoms (LUTS) (1–4). In addition, the widespread prevalence of BPH is a significant U.S. healthcare burden currently estimated at $4 billion per annum with costs expected to increase over the next two decades due to population demographics (4–7). Characteristic histopathological features of BPH include hyperproliferation of the epithelial and stromal prostate compartments resulting in the enlargement of the gland (8–12). The main pharmacological treatment options are currently limited to two approaches: *a)* reducing prostate size by blocking the conversion of testosterone (T) to dihydrotestosterone (DHT) using a 5α-reductase inhibitor (5ARI); and/or, *b)* relaxing the smooth muscle tone within the prostate and bladder neck by using an alpha-adrenergic receptor blocker (α-blocker) (13). However, as the Medical Therapy of Prostatic Symptoms (MTOPS) clinical study reported, these treatment options often fail thus leaving surgical treatment as the only remaining option (14, 15). Therefore, there is a clear need to identify and develop molecular-based approaches for the treatment of the disease.

Budding and branching morphogenesis, a key process in prostatic development, has been posited to lead to the manifestation of the hyperplastic phenotype in BPH (16–20). However, the precise molecular trigger(s) that lead to the resurgence in budding and branching morphogenesis in BPH remain elusive. Investigation of the molecular mechanisms responsible for the budding and branching phenotypes could in turn lead to a better understanding of the causative factors of BPH pathogenesis and thereby point to novel and effective therapeutic strategies for the treatment of the disease.

In this study, using clinical BPH samples and 2D/3D cell culture models, we have utilized a combination of bioinformatic-based analyses, genetic-modulation studies, and a proof-of-concept pharmacological approach to identify TIAM1-RAC1 signaling as a: *1)* driver of budding and branching phenotype in BPH, and *2)* promising candidate pathway for molecular-based approach for therapeutic targeting of BPH.

## Methods

### RNA-Sequencing data processing and analysis

Three mRNA expression datasets (21–23) were obtained from the Genotypes and Phenotypes (dbGap) (https://www.ncbi.nlm.nih.gov/gap/) and the Gene Expression Omnibus (GEO) databases (https://www.ncbi.nlm.nih.gov/geo/). **Table S 1** outlines the information regarding the datasets utilized in the current study. Briefly, the first study, Jin et al (23), utilized 30 BPH and 14 control samples for the RNA-seq. The transition zone (TZ) from BPH was collected at the time of bladder outlet obstruction surgery for the treatment of LUTS. The control tissue used was benign TZ tissue from patients that underwent radical prostatectomy for low volume, low grade prostate cancer that was localized to the peripheral zone (PZ). The second study, Liu et al (21), utilized 18 BPH and 4 control samples for the RNA-seq. Prostate tissue from TZ of BPH patients was utilized for the study. The controls for RNA-seq were obtained from men undergoing radical prostatectomy for prostate cancer without BPH. The third study, Middleton et al (22), that utilized 37 (28 were available in dbGap) BPH and 19 (18 were available in dbGap) controls, acquired BPH tissues from radical prostatectomies for the diagnosis of prostate cancer that had concurrent BPH. The control tissue was obtained from normal PZ of the prostates without other pathologic changes.

The DNASTAR SeqMan NGen assembler (DNASTAR 2016, Madison, WI, USA) was used to map the FASTQ files to the human reference genome (GRCh38). The assembled RNA-seq reads were normalized by the RPKM (Reads Per Kilobase per Million mapped reads) method. Differences in gene expression levels were analyzed using Fisher’s Exact Test in the ArrayStar software package (DNASTAR 2016, Madison, WI, USA). Differentially expressed genes (DEGs) were filtered with a fold-change of ≥ |1.5| and False Discovery Rate (FDR) of < 0.05. An online Venn analysis tool (http://bioinformatics.psb.ugent.be/webtools/Venn/) was utilized to overlap the three sets of DEGs in order to arrive at the common DEGs (cDEGs) – i.e., the shared BPH transcriptomic signature. The heatmaps and boxplots were generated using Pheatmap (https://cran.r-project.org/web/packages/pheatmap/index.html) and ggPlot2 (https://cran.r-project.org/web/packages/ggplot2/index.html). The Database for Annotation, Visualization and Integrated Discovery (DAVID) (https://david.ncifcrf.gov/) was used to analyze Gene Ontology (GO) to generate functional annotations for cellular and molecular functions, including cellular components (CC), biological processes (BP) and molecular functions (MF). Reactome (24) analysis was used to determine the biological functions of the cDEGs and putative upstream regulators were determined by Ingenuity Pathway Analysis (IPA) (QIAGEN Inc., https://digitalinsights.qiagen.com/IPA) (25). cDEGs were interrogated for potential therapeutic candidate compounds using Connectivity Map (CMap) (26, 27), a widely used *in silico* drug screening tool that uses transcriptional expression data to investigate relationships between disease conditions and therapeutic modalities on the basis of altered biological processes or pathways.

### Cell Culture

Human benign prostatic hyperplasia cell lines, BPH-1, BHPrE1, BHPrS1, and normal prostate cell line, NHPrE1, were provided by Dr. Simon Hayward, North Shore University Health System Research Institute, IL, Chicago. Normal prostate human epithelial cell line RWPE-1 was purchased from American Type Culture Collection. Each cell line was maintained in a specific media at 37°C in a humidified CO_2_ (5%) incubator. BPH-1 cells were maintained in Roswell Park Memorial Institute Medium (RPMI; Corning, Cat # 10040CV) supplemented with 10% Cosmic Calf Serum (CCS; HyClone, Cat # 16777-244) and 1% penicillin and streptomycin. BHPrE1 cells were maintained in Dulbecco’s Modified Eagle Medium (DMEM F-12; Corning, Cat # 10090CV), 5% Fetal Bovine Serum (FBS; Gibco, Cat # 10437-028), 1% penicillin and 1% streptomycin (Cytiva, Cat # SV30010), 0.4% Bovine Pituitary Extract, (BPE; Gibco, Cat # 13028014), 10 ng/ml Epidermal Growth Factor, (EGF; Millipore, Cat # E4127) and 1% Insulin-Transferrin-Selenium (ITS; Gibco, Cat # 41400045); BHPrS1 cells were maintained in RPMI-1640 with 5% FBS and 1% penicillin and streptomycin; RWPE-1 cells were maintained in keratinocyte medium (Gibco, Cat # 10724011) containing BPE and EGF. The cell lines used for this study were intermittently evaluated in-house for mycoplasma contamination. NSC23766 was purchased from Tocris (Cat # 21-611-0R) with the purity of at least 96%. NSC23766 was reconstituted in molecular grade water in concentration of 100 mM and kept in −80 °C for further experiments.

### RNA isolation, cDNA synthesis and quantitative real-time RT-PCR (qRT-PCR)

Total RNAs were extracted from human benign prostatic cells using RNeasy Mini Kit as per manufacture’s protocol (Qiagen, Cat # 74104). cDNA was synthesized using 2µg of RNA using a high-capacity RNA to cDNA kit (applied biosystems, Cat # 4368814). qRT-PCR was performed using PowerUp™ SYBR™ Green Master Mix (Life Technologies, Cat # A25742) with specific gene primers synthesized from Integrated DNA Technologies (IDT, Coralville, IA). The qRT-PCR analysis was performed on the QuantStudio™ 12K Flex platform.

### Cell lysate preparation and Western blot analysis

Total protein lysates were prepared from the cells followed by protein quantification using the protein DC assay kit (Bio-Rad, Cat # 5000-001) as described previously (28). Extracted proteins were resolved on a 10% SDS-PAGE gel and electroblotted onto a PVDF membrane (Sigma Aldrich, Cat # IPVH00005). The membranes were blocked with 1X Blocker^TM^ FL 10X Fluorescent blocking buffer (Thermo Fisher Scientific, Cat # 37565) followed by incubation with the respective primary antibodies overnight at 4°C. The detailed list of the antibodies used in this study is provided in the supplementary information **(Table S 2)**. Subsequent to washing with TBST (0.1% Tween), the membranes were probed with HRP-conjugated secondary antibodies at 1:2000 dilution (rabbit/mouse, Cell Signaling Technology) for 1 hour at RT. The bands were visualized using west Pico chemiluminescent kit (Thermo Fisher Scientific, Cat # 34580) under Chemi-doc touch imaging system (Bio-Rad, CA, USA). β-Actin was used as the loading control.

### Immunohistochemistry of human samples

After obtaining IRB approval (IRB # L22-235), immunohistochemical (IHC) analysis was performed on de-identified, paraffin-embedded human prostate tissues: BPH (n=14) and control tissues (n=10). To reduce batch effects, all samples were processed together for IHC analysis. In brief, slides were incubated in xylene (Fisher chemical, Cat # X3S-4) three times (5 min each) for deparaffinization. Following a series of 5 min incubations in ethanol at decreasing concentrations (100%–10%) and three phosphate-buffered saline (PBS) washes for 5 min each, slide sections were hydrated by rinsing under running water. Using 1X antigen unmasking buffer (Vector Lab, Burlingame, CA, USA, Cat # H-3300), antigen unmasking was carried out in a decloaking chamber (Oster). This was followed by blocking of endogenous peroxidases using Bloxall (Vector Lab, Cat # SP-6000). Next, Avidin and Biotin blocking (Vector Lab, Cat # SP-2001) was performed followed by a PBS wash. Goat serum (Vector Lab, Cat # S-1000) at a dilution of 5% in PBS was used to block the tissue section. Next, the slides were incubated overnight with Anti-TIAM1 primary antibody (Abcam, Cat # ab211518, 1:800 dilution) at 4 °C in a humidified chamber. The following day, three PBS washes were performed for 5 min each and the slides were incubated for 1 hour with the secondary antibody (Vector Lab, BA-1000, 1:200 dilution), followed by three PBS washes for 5 min each. The slides were then incubated with premixed avidin biotin complex (Vector Lab, Cat # PK-6101) for 45 min and then washed twice with PBS. For the development of color, peroxidase substrate (Vector Lab, Cat # SK-4105) was added, followed by washing the slides with water, counterstaining using hematoxylin (Vector Lab, Cat # H-3401) and then rinsing again with water before mounting with Vectamount (Vector Lab, Cat # H-5000).

### Image acquisition and analysis

The IHC stained slides were analyzed and scored based on the immunoreactive score (IRS) (29). In short, IRS which is used to quantify the IHC staining results by evaluating the percentage of positive cells and the intensity of staining. For this analysis, the pathologist visually estimated the proportion of cells showing immunoreactivity (positive cells) and assessed the strength of the staining intensity. The IHC score was calculated as IRS = (percentage of positive cells) x (intensity of staining). The images were acquired using Olympus XI83 microscope.

### Proliferation assay

Cell proliferation was measured using WST-1 (water-soluble tetrazolium) assay (Roche, Cat # 501003295) according to standard manufacturer’s protocol. 2000 cells per well were seeded in a 96-well plate with 100 μL of culture medium and incubated at 5% CO2, 37 °C overnight. The following day, differing concentrations of NSC23766, ranging from 1.25 μM to 200 μM or vehicle control (molecular grade water) were administered and the plates were incubated for 5 days. Cell proliferation was examined at 24, 48, 72, 96-and 120-hours using WST-1 reagent. Absorbance was measured at 450 nm using an iMark microplate reader (Bio-Rad).

### Organoid assay

For the mono-culture organoid assay, 700 cells/well were mixed in 300 μL of 5 μg/ml of Matrigel (Corning; Cat # CB40230) in the media and seeded on a 150 μL of 5 μg/ml Matrigel base layer. The assay was performed as per the manufacturer’s protocol, Corning LifeSciences (30) and previously published protocols (19, 31–33). The cultures were maintained in 500 μL of media that was changed every 2-3 days. For co-culture experiments, 700 epithelial cells were seeded in Matrigel along with 1400 stromal cells. The co-cultures were maintained in 500 μL media and media was changed every 2-3 days. The organoids were analyzed for size, number, buds and branches using the inbuilt EIS microscope software (Nikon microscope) and Image J (34).

### Lentiviral vector

HEK293T cells (1.2×10^7^) were co-transfected with either lentiviral vector containing constructs, pLKO.1-Neo-CMV-tGFP TIAM1 shRNA plasmid (Sigma Aldrich, Cat # 07202334MN, TRCN0000256946 that targets “TTCGAAGGCTGTACGTGAATA”) or pLKO.1-Neo-CMV-tGFP-Scramble shRNA plasmid (Addgene#136035 as scramble control). Lentiviral vectors were packaged using packaging plasmid (pCMV delta R8.2 dvpr, Addgene #8455) and the envelope plasmid (pCMV-VSV-G, Addgene #8454). After 72 hours of transfection, the virus containing media was collected and filtered and used for the BHPrE1 cells. After three days, cells were treated with 300 µg/ml neomycin to obtain stable TIAM1 knock-down clones.

### Statistics

Statistical analyses were performed using GraphPad Prism version 9.0.1 (GraphPad Software, San Diego, CA). The unpaired two-tailed student’s t-test was employed for comparing two groups, one-way ANOVA, and two-way ANOVA were utilized to determine statistical significance for three or more groups. Differences were considered statistically significant when *p < 0.05. All the experiments were done N=3 and data are presented as mean ± standard deviation (S.D.).

## Results

### Identification of a BPH transcriptomic signature

In the current study, mRNA expression profiles from three independent BPH patient datasets were utilized to compare mRNA expression between BPH and control tissues. Through comparing the RNA-Seq read counts of various genes followed by the application of the cutoff criteria (fold-change ≥|1.5| and FDR of <0.05), DEGs were identified for each dataset. Next, the DEGs obtained from each of the three datasets were overlapped to arrive at 84 cDEGs which constituted the putative BPH transcriptomic signature (**Fig. 1 A**).

**Fig. 1:**
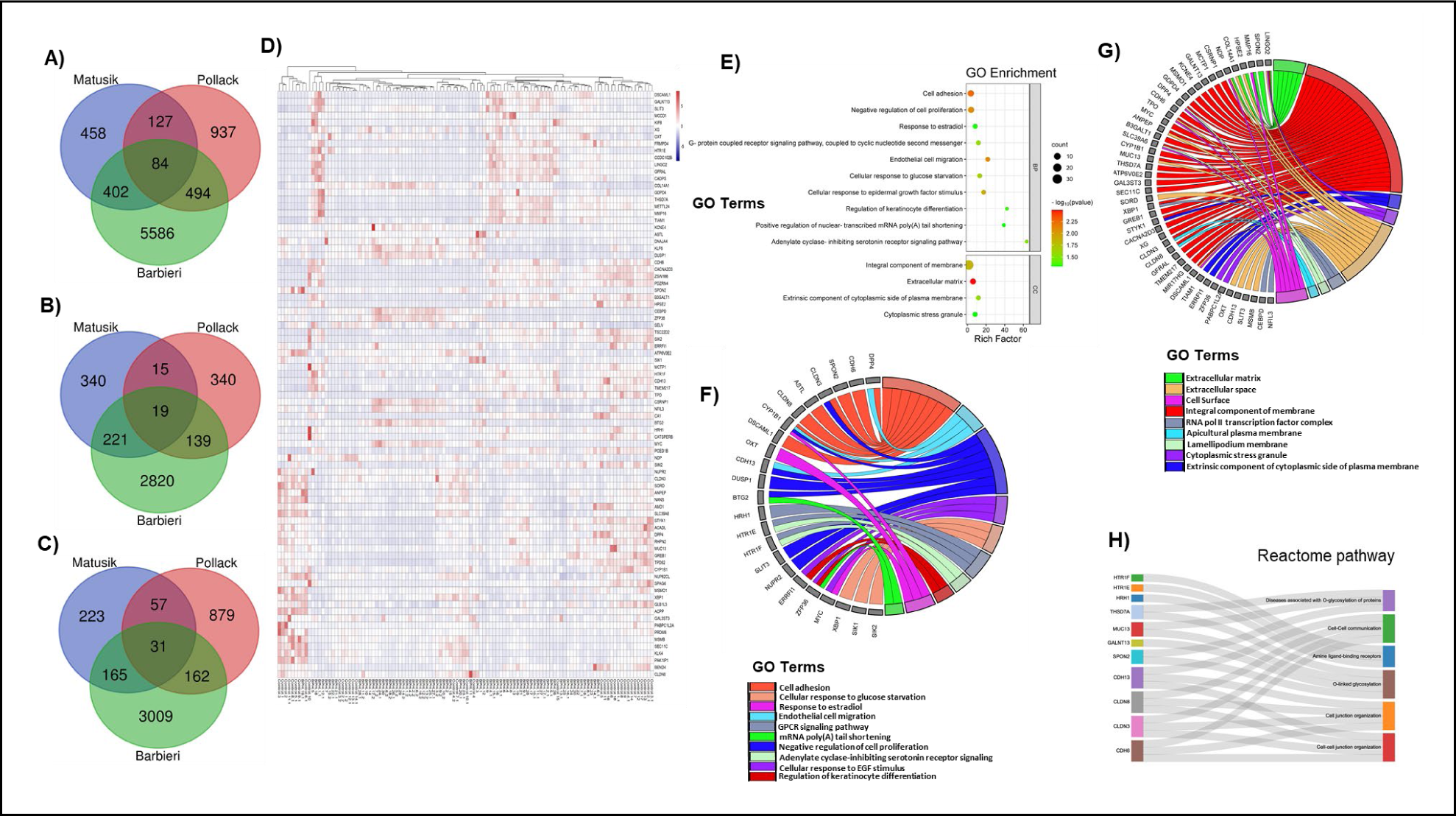
Identification of a BPH transcriptomic signature. **A)** Venn diagram showing 84 common differentially expressed genes (cDEGs) based of the overlap of individual sets of DEGs obtained from RNA-Seq datasets from three independent BPH patient cohorts (Matusik, Pollack, Barbieri). (Differentially expressed genes (DEGs) were filtered with a fold-change of ≥ |1.5| and False Discovery Rate (FDR) of < 0.05). **B)** Venn diagram showing 19 upregulated cDEGs from the three datasets. **C)** Venn diagram showing 31 downregulated cDEGs from the three datasets. **D)** Heatmap representing the normalized patient-wise expression pattern of the cDEGs from the three datasets. **E)** Gene ontology (GO) pathway analysis of the 84 cDEGs in two functional groups: biological processes (BP) and cell compartment (CC) (p < 0.05). **F)** Circular genome visualization (Circos) plot for illustrating the correlation between biological processes and their associated cDEGs (p< 0.05). **G)** Circos plot illustrating the correlation between cellular compartments and their associated cDEGs. **H)** Reactome pathway analysis showing the significantly enriched Reactome pathways from the 84 cDEGs.

The cDEGs included 19 cDEGs consistently up-regulated among all three datasets (**Fig. 1 B**), and 31 cDEGs consistently downregulated in all three datasets (**Fig. 1 C**). The remaining 34 cDEGs were significantly dysregulated but not consistently up-or down-regulated in all three datasets. A heatmap of the 84 cDEGs was created (**Fig. 1 D**). To garner insights regarding the possible biological functions of the cDEGs, gene enrichment analysis was performed using Database for Annotation, Visualization and Integrated Discovery (DAVID), which included gene ontology (GO) **(Fig. 1 E-G)** and Reactome pathway enrichment analyses **(Fig. 1 H)**. For GO biological processes (BP), cell adhesion, negative regulation of cell proliferation, response to estradiol, differentiation and migration were implicated (**Fig. 1 E, F**). For the GO cellular component (CC), gene enrichment primarily involved membrane and extracellular matrix (**Fig. 1 E, G**). Functional implications of the cDEGs were further investigated by Reactome pathway analysis. A number of cDEGs were enriched in six Reactome pathways, including glycosylation of proteins and cell-cell communication **(Fig. 1 H)**. Taken together, a BPH transcriptomic signature was identified that is associated with key biological BPH processes including cell-cell communication, cellular proliferation, and adhesion.

### Identification of putative upstream regulators of BPH transcriptomic signature

Gene expression data and transcriptomic signatures (**Fig. 2 A**) have frequently been used as diagnostic and prognostic markers of disease. However, gene expression signatures often include several passenger genes that may not represent disease drivers. On the other hand, upstream regulators often represent crucial factors that drive progression and disease etiology and are better candidates for potential therapeutic interventions. Therefore, QIAGEN Ingenuity Pathway Analysis (IPA) (25) was performed to identify key upstream regulators of the 84 cDEGs. The top five upstream regulators (p-value < 0.05) are listed in **Fig. 2 B** include G protein-coupled estrogen receptor 1 (GPER1), KLF transcription factor 5 (KLF5), CTR9 Homolog, Paf1/RNA Polymerase II Complex Component (CTR9), mir-let7 (let-7), and Rac family small GTPase 1 (RAC1).

**Fig. 2:**
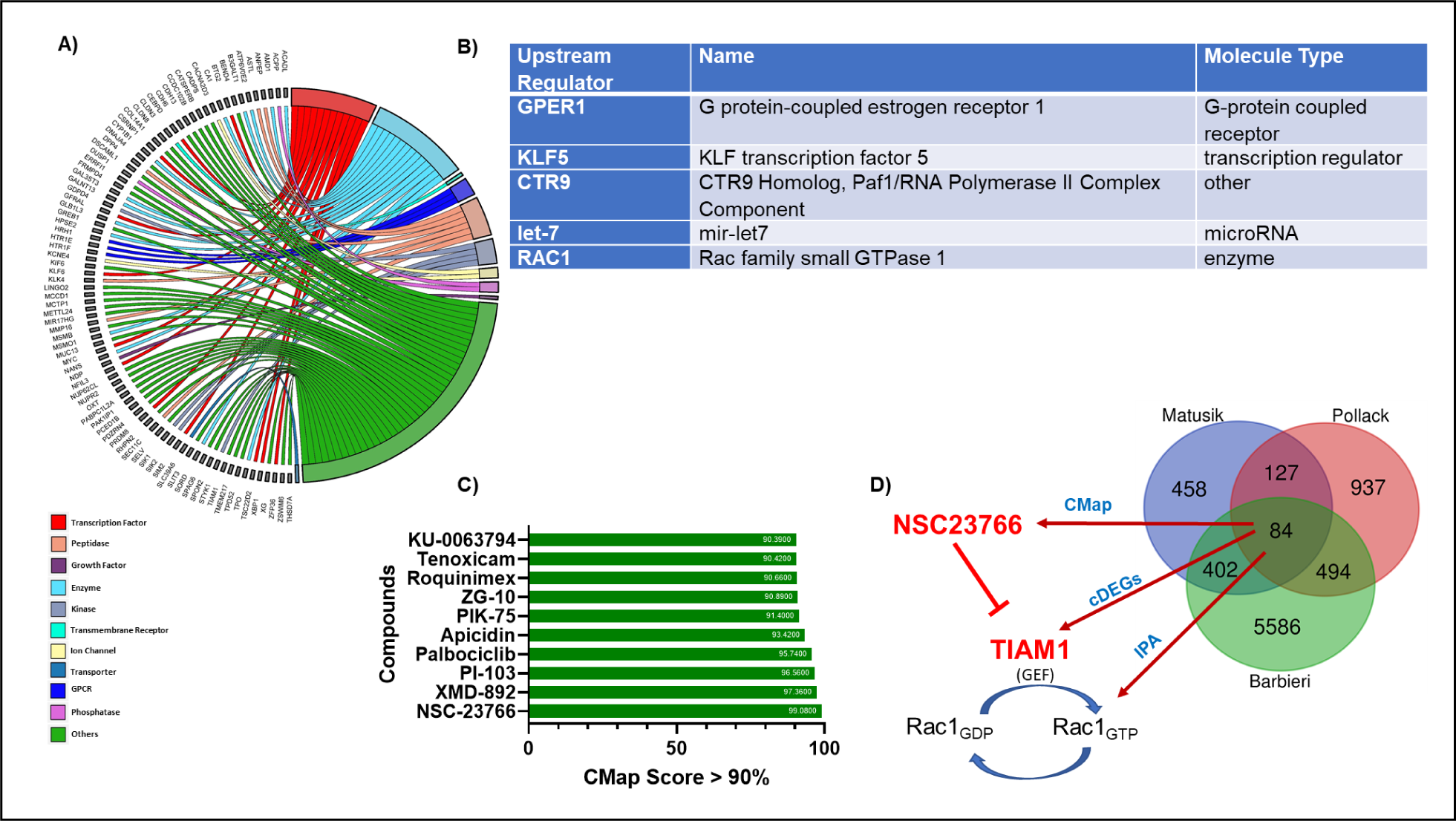
Identification of the upstream regulators and potential therapeutic compounds drug candidates based on the BPH transcriptomic signature. **A)** Circos plot illustrating the correlation between cDEGs and their functional categories: transcription factors, peptidases, growth factors, enzymes, kinases, transmembrane receptors, ion channels, transporters, G-protein coupled receptors, and phosphatases. **B)** Ingenuity Pathway Analysis (IPA) showing key upstream regulators of the 84 cDEGs. **C)** Connectivity Map (CMap) analysis showing the CMap scores of the top 10 nominated compounds based on the 84 cDEGs. **D)** A schematic showing the connecting links between: 1) TIAM1 (a RAC1 specific GEF) as one of the upregulated cDEGs, 2) RAC1 as an upstream regulator based on the 84 cDEGs, and 3) NSC23766, a small molecule inhibitor of TIAM1-RAC1 signaling, as the top candidate compound from the CMap analysis.

### Identification of NSC23766 as a potential inhibitor impacting the BPH transcriptomic signature

The cDEGs were queried to discover potential pathway inhibitors using Connectivity Map (CMap), (L1000 platform; https://clue.io/l1000-query) a widely used *in silico* drug screening tool that uses cellular responses to perturbation to illustrate relationships between genes, diseases, and therapeutic compounds through connectivity score (CMap score) (26, 27). CMap score is a measure of the degree of correlation with the gene expression signature. Using CMap analysis, the connectivity scores were calculated for both upregulated and downregulated cDEGs and therapeutic compounds with a CMap score of 90 or higher were shortlisted as potentially impacting key aspects of the BPH signature **(Fig. 2 C and Table S 3)**. Interestingly, the shortlisted compounds included inhibitors of inflammation (8, 9, 12), fibrosis (12), and proliferation (35–37) - the hallmarks of BPH. CMap analysis revealed NSC23766 as the highest ranked candidate compound with a connectivity score of 99.08% **(Fig. 2 C).** NSC23766 is a small molecule that blocks activation of Rac family small GTPase 1 (RAC1) protein by inhibiting T-lymphoma invasion and metastasis-inducing protein-1 (TIAM1) activity (38, 39) **(Fig. 2 D)**; the list of upstream cDEGs regulators included the GTPase, RAC1 **(Fig. 2 B)**.

### TIAM1 protein expression is increased in BPH

Since: *1)* NSC23766 is a small molecule inhibitor of TIAM1 activity (39); *2)* both, Ingenuity Pathway Analysis (IPA) and Connectivity Map (CMap) analysis, pointed to TIAM1-RAC1 signaling as the potential lead pathway for therapeutic targeting of BPH **(Fig. 2 D)**; and *3)* our BPH transcriptomic signature revealed TIAM1 upregulation in all three datasets - we sought to validate increased expression of TIAM1 in an independent fourth cohort of BPH patients. We therefore probed for expression of TIAM1 protein in human prostate control **(Fig. 3 A)** and BPH **(Fig. 3 B)** specimens using immunohistochemistry (IHC) **(Figs. 3 A-C, Fig. S 1)**. BPH specimens are from the transition zone tissues of patients who had undergone simple prostatectomy. Control specimens are benign transition zone tissues from patients who had undergone radical prostatectomy for low-volume prostate cancer that was localized in the peripheral zone. The results revealed a significant increase in TIAM1 expression in BPH tissues compared to control tissues **(Fig. 3 C)**. Taken together, the differential increase of TIAM1 protein compared to control tissues was consistent with the differential increase of mRNA levels **(Fig. 1 D)**.

**Fig. 3:**
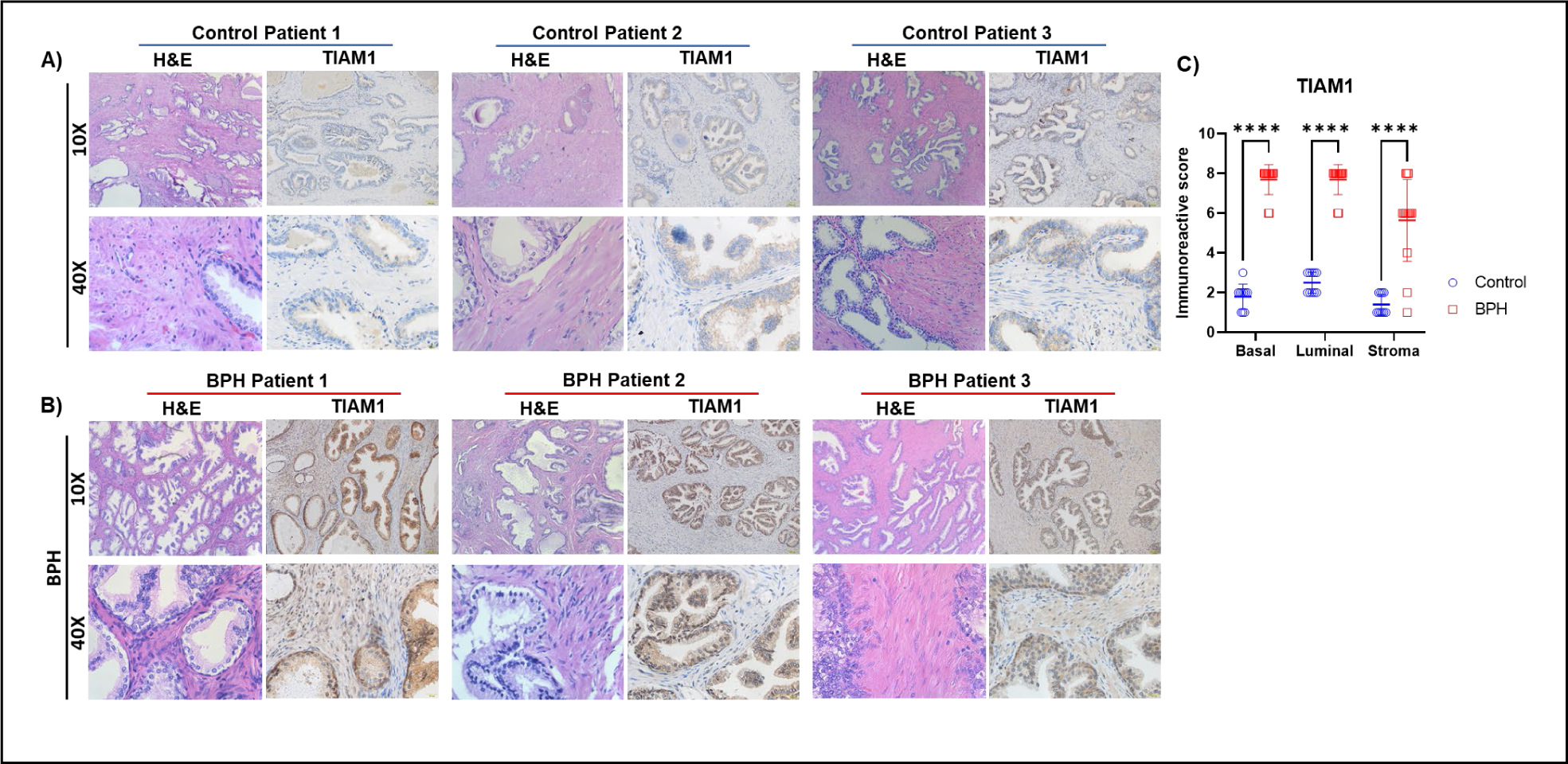
TIAM1 protein expression is increased in human BPH tissues. **A, B)** Representative IHC and H&E staining showing TIAM1 expression at 10x and 40x magnification in control (A) and BPH tissues (B). **C)** Quantification of the IHC images using the immunoreactive score (IRS) scores in BPH (n=14) compared with control (n=10). The immunoreactive score (IRS)= % of cells X intensity of staining. Two-way ANOVA. pvalue: ****<0.0001.

### NSC23766 caused a decrease in the proliferation of human benign prostatic epithelial and stromal cells

Increased proliferation of both epithelial and stromal cells is a defining feature of BPH. Therefore, to explore the effects of NSC23766 on the proliferation of BPH cells, we treated human benign prostatic epithelial cell lines, BHPrE1, BPH-1, NHPrE1 and RWPE-1 and human prostatic stromal cells, BHPrS1, with concentrations of NSC23766 ranging from 1.25 µM to 200 µM. After 24 hours of NSC23766 exposure, a significant decrease in proliferation was observed in the NSC23766 treated cells in a dose-dependent manner across the epithelial and stromal cell lines **(Fig. 4 A-B)**.

**Fig. 4:**
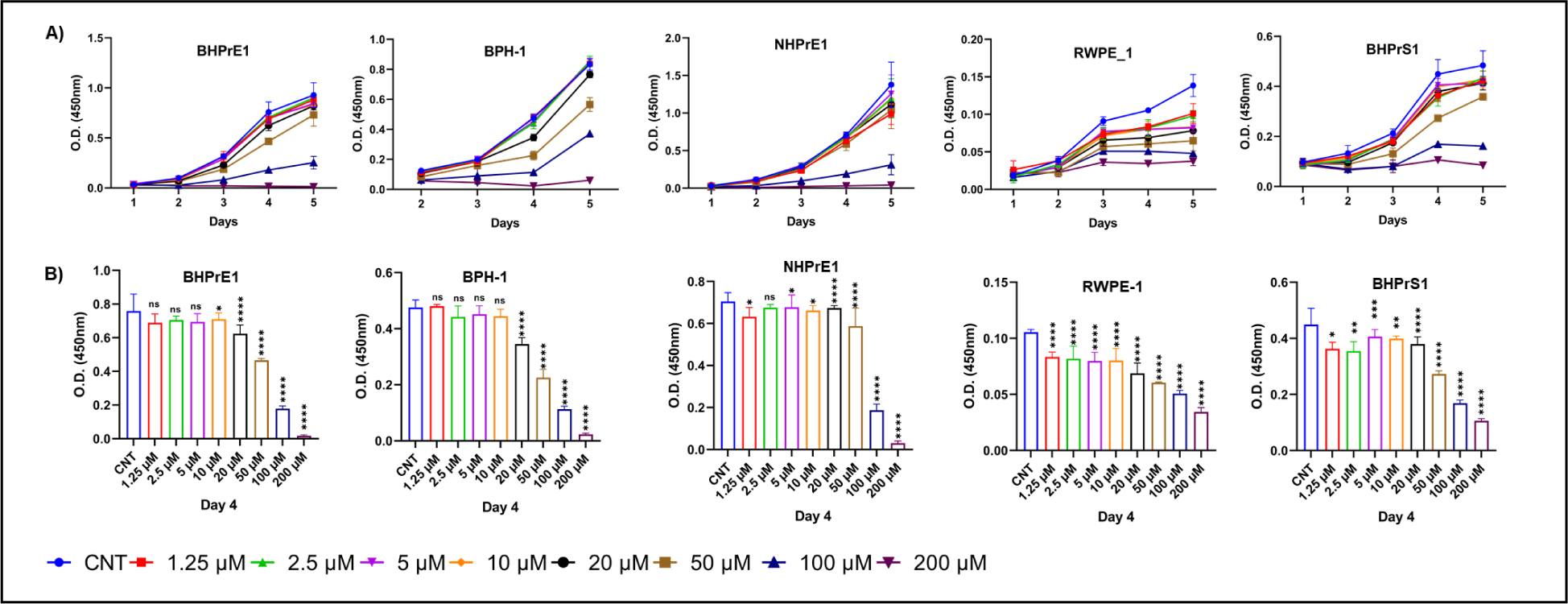
NSC23766 caused a decrease in the proliferation of human benign prostatic epithelial and stromal cells. **A)** WST-1 assay-based cell proliferation analysis of benign prostate cell lines, following 5 days treatment with NSC23766 at concentrations ranging from 1.25 µM to 200 µM. **B)** Quantification of different concentration of NSC23766 treatment on day 4 in deferent benign prostate cell lines. (N=5 in each condition, Mean ± standard deviation *, p < 0.05; **, p < 0.01; ***, p< 0.001; ****; p < 0.0001).

### NSC23766 caused a decrease in the budding and branching phenotypes in organoids from human benign prostatic epithelial cells

Budding and branching, which lead to new glandular structures, are phenotypes associated with BPH. To test the effect of NSC23766 on BPH budding and branching phenotypes, 3D-organoid-culture model of human benign prostatic cell lines was employed. Organoids were formed from BHPrE1, NHPrE1 and RWPE-1 human benign prostatic epithelial cell lines; cells organize into basal and luminal layers and eventually bud and branch to form prostatic glands in 3D culture (19, 40, 41). The BHPrE1 and RWPE-1 organoids were stained using immunofluorescence (IF) staining for keratin 5 (KRT 5) and keratin 8/18 (KRT8/18) which stain basal and luminal epithelial cells, respectively (**Fig. 5 A).** The protein expression of TIAM1 in the BHPrE1 organoids was also visualized using IF staining (**Fig. S 2)**. The effect of NSC23766 on the budding and branching of BHPrE1organoids was then tested. NSC23766 treatment starting from Day 1 of the organoid cultures resulted in a decrease in the number of budded and branched organoids in a dose-dependent manner, p< 0.05 and p<0.0006 (**Fig. 5 B-E**).

**Fig. 5:**
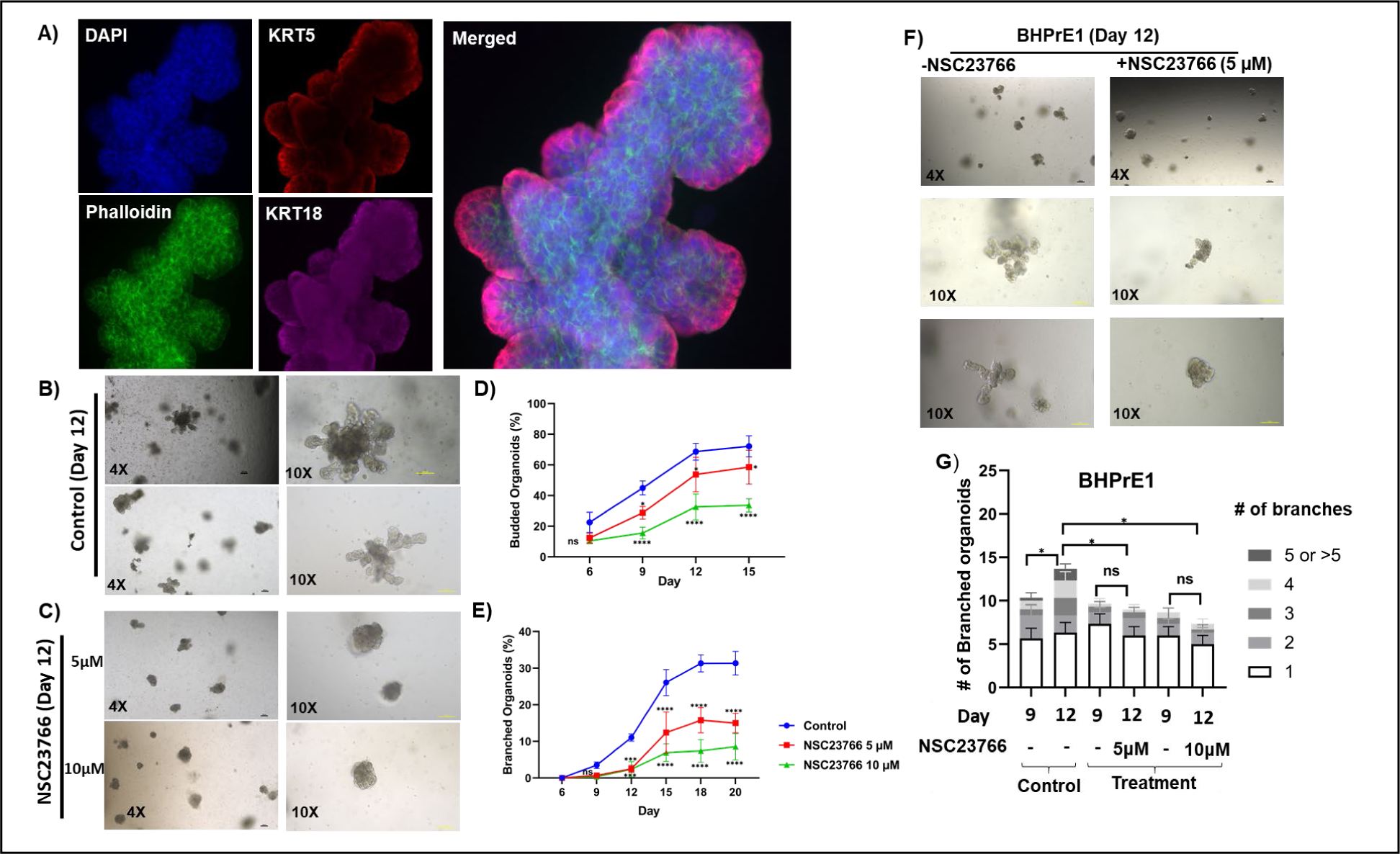
NSC23766 caused a decrease in the budding and branching phenotypes in organoids from human benign prostatic epithelial cells. **A)** Whole-mount Immunofluorescence staining of BHPrE1 organoids with basal cell marker (KRT5) and luminal cell marker (KRT8/18) on day 9. **B)** Representative images taken on day 12 of BHPrE1 organoids. **C)** Representative images taken on day 12 of BHPrE1 organoids with NSC23766 treatment starting on day 1 at 5 µM and 10 µM concentrations. **D)** Quantification of the percentage of BHPrE1 budded organoids with NSC23766 treatment starting on day 1 at 5 µM and 10 µM concentrations compared with vehicle control. **E)** Quantification of the percentage of BHPrE1 branched organoids with NSC23766 treatment starting on day 1 at 5 µM and 10 µM concentrations compared with vehicle control. **F)** Representative images taken on day 12 of BHPrE1 organoids with NSC23766 treatment starting on day 9 at 5μM concentration or vehicle control. **G)** Quantification of the percentage of BHPrE1 branched organoids on day 9 and day 12, following treatment with NSC23766 on day 9. (N=3, Mean ± SD, two-way ANOVA was performed, *, p < 0.05; **, p < 0.01; ***, p< 0.001; ****; p < 0.0001).

In another experiment, the effect of NSC23766 on BHPrE1 organoids with pre-existing branches was evaluated. For this purpose, BHPrE1 organoid cultures were exposed to NSC23766 starting on Day 9, a time-point at which most of organoids had begun forming branches - and branches were re-evaluated on Day 12. The number of branched organoids in the vehicle control continued to increase from Day 9 to Day 12, however, the number of branched organoids did not change significantly in the treatment group **(Fig. 5 F, G)**. The total number of branched organoids on Day 12 in the NSC23766 treatment groups was also decreased when compared with the vehicle control on Day 12 (p<0.02; **Fig. 5 F, G)**.

To test if discontinuation of NSC23766 treatment could reinitiate the organoid budding and branching, treatment was started on Day 9 when most of the organoids had begun forming branches and then discontinued the treatment for the next 3 days in order to test if the budding and branching phenotypes could be reinitiated. While continued treatment with NSC23766 (from Day 9 to Day 16) caused a decrease in the percentage of organoids with buds and branches; however, discontinued NSC23766 treatment from day 12 resulted in re-initiation of budding and branching (**Fig. S 3**).

We observed similar effects of NSC23766 on RWPE-1 organoids Treatment of organoids either from Day 1 or Day 9 resulted in decreases of size and number of branched organoids in a dose-dependent manner (p< 0.04; **Fig. S 4 A-G**). Similar results were observed with NHPrE1 organoids (**Fig. S 5**). Taken together, results indicated that NSC23766 treatment resulted in the reduction of budding and branching phenotypes of organoids formed from multiple benign prostatic epithelial cells in a dose-dependent manner.

### NSC23766 caused a decrease in budding and branching phenotypes in organoids from human benign prostatic epithelial cells in the presence of stroma

Since prostatic stroma has an established role in glandular differentiation of the prostate and pathophysiology of BPH (18, 42, 43), we evaluated the effects of NSC23766 on the budding and branching of BHPrE1 organoids when co-cultured with BHPrS1 human prostate stromal cells (**Fig. 6 A**). Results demonstrated that co-culture with stromal cells significantly increased the number of BHPrE1 organoids formed, p-value: <0.0001 **(Fig. 6 B)**. Further, co-culture significantly enhanced the branching efficiency of BHPrE1 organoids with an observed increase in the number of organoids with multiple branches **(Fig. 6 C)**.

**Fig. 6:**
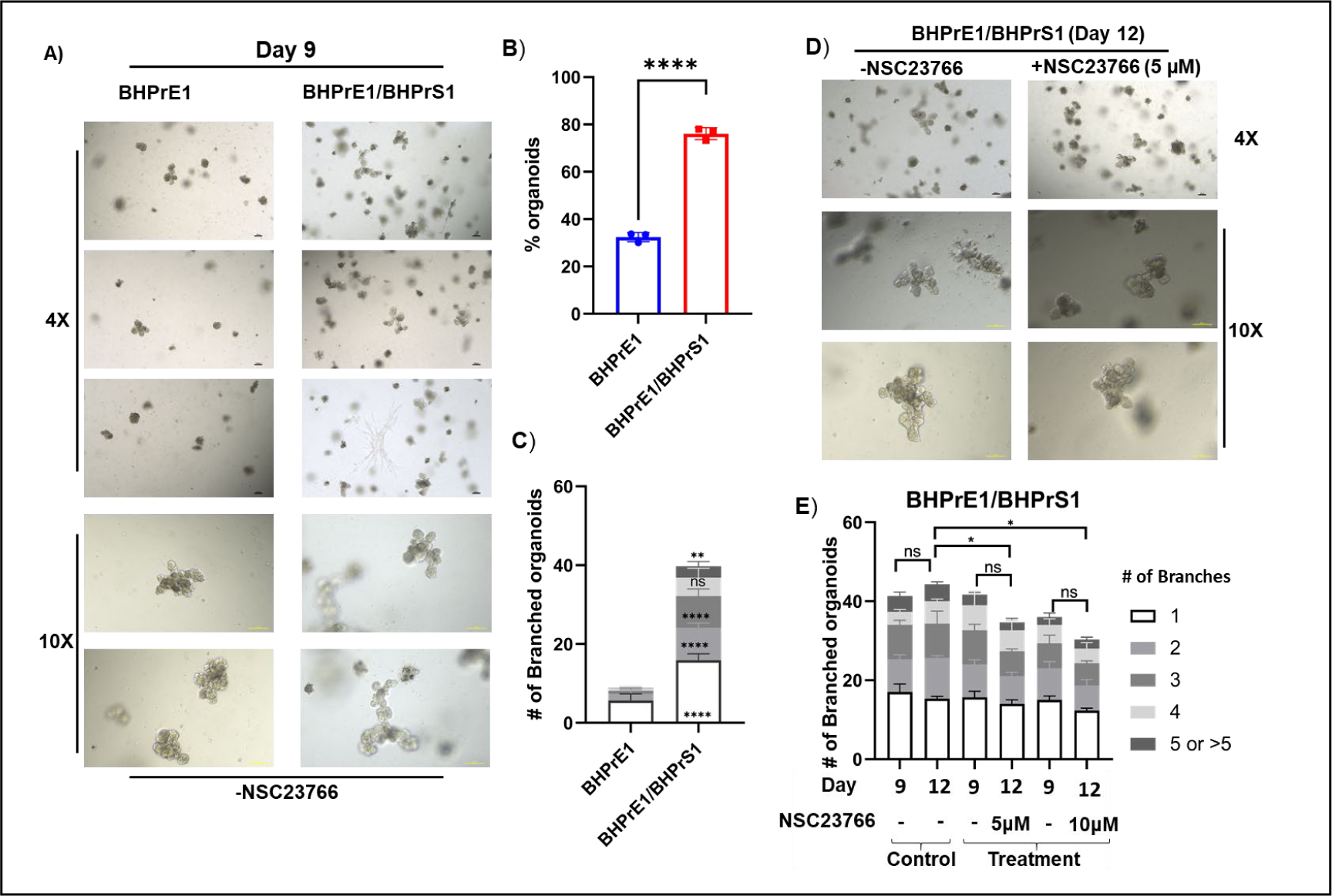
NSC23766 caused a decrease in budding and branching phenotypes in organoids from human benign prostatic epithelial cells in the presence of stroma. **A)** Representative images taken on day 9 of BHPrE1 organoids in mono-culture and co-culture with stroma. **B)** Quantification of the percentage of BHPrE1 organoids in mono-culture and co-culture with BHPrS1 cells. **C)** Quantification of the number of branched organoids with multiple of branches ranging from 1 to 5 branches in mono-culture and co-culture with stroma on day 9. **D)** Representative images taken on day 12 of BHPrE1 organoids in co-culture with stroma treated with 5μM NSC23766 or vehicle control. **E)** Quantification of the number of BHPrE1 branched organoids in co-culture with BHPrS1 cells on day 9 and day 12, following 5μM NSC23766 treatment starting on day 9. N=3 in each condition, mean ± SD. Student paired t-test and two-way ANOVA was performed, *, p < 0.05; **, p < 0.01; ***, p< 0.001; ****; p < 0.0001.

Next, the effect of NSC23766 on BHPrE1 organoids co-cultured with stromal cells was evaluated. NSC23766 was added on Day 9, and effects assayed on Day 12. In controls, by Day 12 there was an increase in the number of branched BHPrE1 organoids. However, NSC23766 treatment prevented the increase. (**Figs. 6 D, E**). Further, in the stromal co-culture conditions, the total number of branched BHPrE1 organoids on Day 12 in the treatment group was significantly reduced when compared with the vehicle control on Day 12 (p<0.02; **Figs. 6 D, E)**. These results suggest that NSC23766-mediated phenotypes of attenuated prostatic budding and branching is maintained in the physiological context of the presence of stroma.

### Genetic knock-down of TIAM1 in human benign prostatic epithelial cells phenocopies the effects observed due to NSC23766 treatment

To test if NSC23766, a small molecule inhibitor of TIAM1, phenocopies the genetic knock-down of TIAM1 in human benign prostatic cells, we performed an sh-RNA-based knock-down of TIAM1 in BHPrE1 cells. Following validation of TIAM1 knock-down **(Fig. 7 A, B and C)**, cell proliferation analysis revealed a significant reduction in cell growth in BHPrE1^shTIAM1^ cells compared to the control BHPrE1^NTSCR^ cells (**Fig. 7 D**). Further, compared to the BHPrE1^NTSCR^ group, organoids formed from BHPrE1^shTIAM1^ cells showed both a significant decrease in the: *1)* total number of branched organoids, and *2)* organoids with 4 or more branches **(Fig. 7 E and F).**

**Fig. 7:**
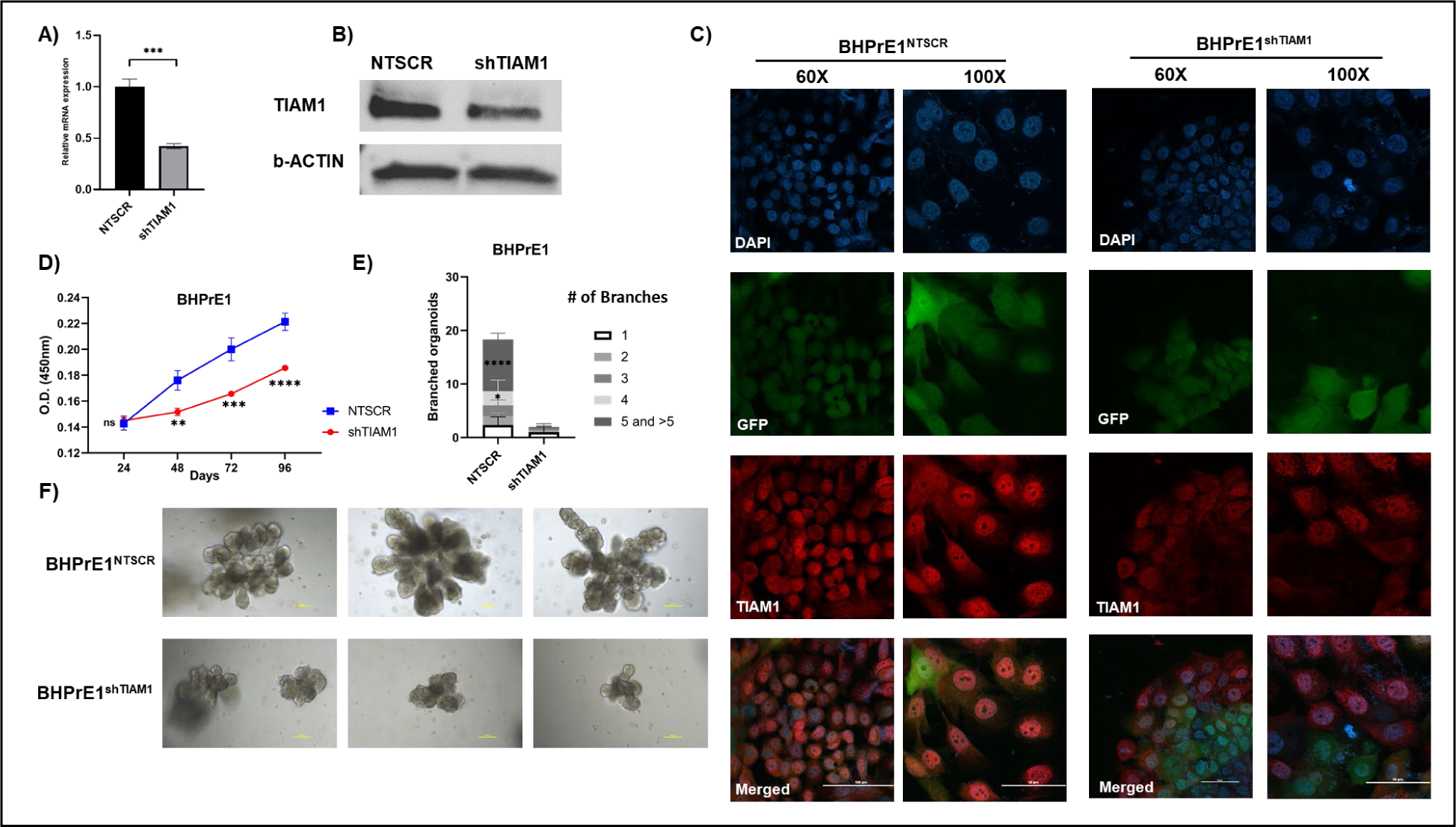
Genetic knock-down of TIAM1 leads to decreases in proliferation of BHPrE1 cells and branching phenotype of BHPrE1 organoids. **A)** qRT-PCR analysis showing decreased mRNA expression of TIAM1 upon shRNA mediated knock-down of TIAM1 in BHPrE1 cells compared with NTSCR control. **B)** Western blot analysis showing decreased protein expression of TIAM1 upon shRNA mediated knock-down of BHPrE1^shTIAM1^ cells compared with BHPrE1^NTSCR^. **C)** Immunofluorescence analysis showing decreased protein expression of TIAM1 upon shRNA mediated knock-down of BHPrE1^shTIAM1^cells compared with BHPrE1^NTSCR^. **D)** WST-1 assay-based cell proliferation analysis of BHPrE1^shTIAM1^ and BHPrE1 ^NTSCR^, showing a decrease in cell proliferation in BHPrE1 due to TIAM1 knock-down. N=5, Mean ± SD. Student paired t-test was performed, **, p < 0.01; ***, p< 0.001; ****; p < 0.0001. **E)** Quantification of the number of BHPrE1^shTIAM1^ branched organoids compared with BHPrE1^NTSCR^ on day 12. N=3, Mean ± SD. One-way ANOVA was performed, *, p < 0.05; ****; p < 0.0001. **F)** Representative images taken on day 12 of BHPrE1^shTIAM1^ organoids compared with BHPrE1^NTSCR^.

## Discussion

In this study, multiple lines of evidence obtained from both bioinformatic and experimental analyses of human BPH pointed to TIAM1-RAC1 signaling as the potential lead candidate pathway for a molecular-based strategy for therapeutic targeting of BPH. *1)* TIAM1 was found to be over-expressed in the transcriptomic signature derived from bioinformatic analysis of transcriptomic datasets from three independent BPH patient cohorts, which was subsequently validated at the protein level in a 4^th^ patient cohort. *2)* Both Ingenuity Pathway Analysis (IPA) and Connectivity Map (CMap) analysis of the transcriptomic signature pointed to TIAM1-RAC1 signaling as the potential lead pathway for therapeutic targeting of BPH. *3)* A proof-of-concept pharmacological approach of TIAM1-RAC1 inhibition using NSC23766 also resulted in attenuated budding and branching of human prostatic organoids in mono-culture and stromal co-culture conditions. *4)* shRNA-based genetic knock-down of TIAM1 in human prostatic cells led to a reduction in budding and branching phenotypes. Therefore, the initial identification of TIAM1 over-expression in BPH through bioinformatic analyses was followed by experimental studies of TIAM1 loss-of-function and pharmacological-based TIAM1-RAC1 inhibition that complemented each other in delineating the biological functions of TIAM1 in BPH **(Fig. 8)**.

**Fig. 8:**
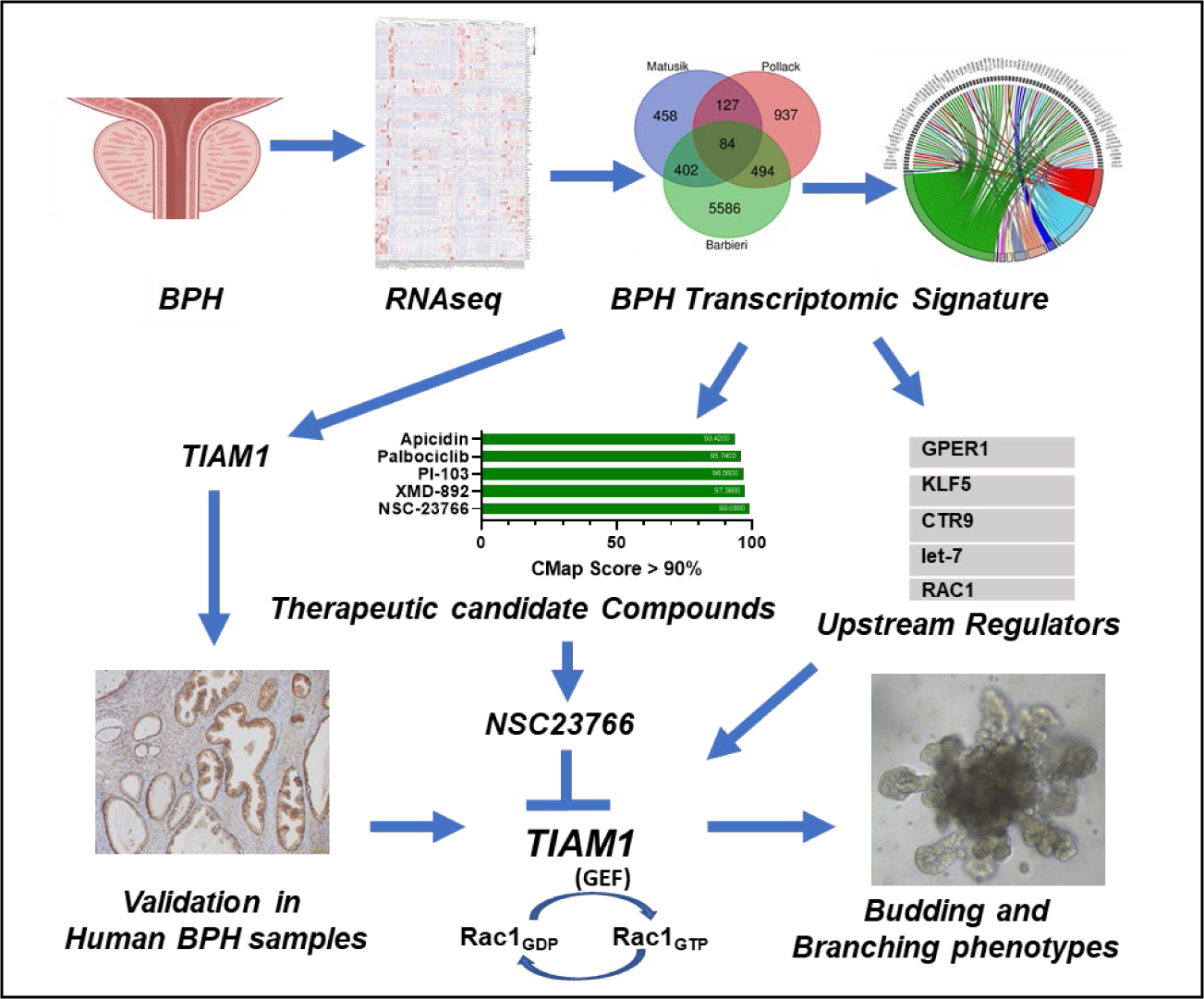
Study workflow that led to the identification of TIAM1 as a potential therapeutic target for BPH: A BPH transcriptomic signature was obtained by the overlap of RNA-seq datasets from three independent BPH patient cohorts. Bioinformatic and experimental analyses based on the BPH transcriptomic signature led to the identification of TIAM1 overexpression in BPH, and NSC23766 - a small molecule inhibitor of TIAM1 signaling - as the lead compound for BPH therapy. Immunohistochemistry validated TIAM1 overexpression in a 4^th^ BPH patient cohort, and exposure of human BPH epithelial and stromal cells to NSC23766 significantly decreased their proliferation and reduced the budding and branching phenotypes of prostatic organoids.

The three datasets used in this study: *1)* belonged to patients with varying past medical histories, *2)* had variances in the control tissue used [Transition Zone (TZ) versus Peripheral Zone (PZ)] (44) and, *3)* incorporated variances in tissue processing post-harvest (RNA isolation from paraffin blocks or frozen tissue). Despite these variances, integration of these datasets not only facilitated the examination of differential gene expression in a greater number of BPH samples, but remarkably revealed a set of cDEGs, a majority of which have previously been reported to be dysregulated in BPH (45–75).

It has been hypothesized that reactivation of budding and branching morphogenesis, a developmental arborization program in the prostate and other organs, drives the BPH hyperplasia phenotype (16, 17). However, the molecular trigger(s) leading to the reactivation of budding and branching morphogenesis in BPH remains elusive. Whereas previous studies focused on prostate ontology have utilized rat and mouse models (76–87), recent reports on prostate pathology have used organoid models (18, 19). Therefore, we investigated the effect of TIAM1 inhibition on prostatic branching by utilizing human benign prostatic epithelial cells that form organoids. These epithelial cells organize into basal and luminal layers, express androgen receptor (AR) and AR-target genes, and eventually bud and branch to form prostatic glands in 3D culture (18). Our results revealed that administration of NSC23766 resulted in attenuated budding and branching of organoids formed from BHPrE1 prostatic epithelial cells.

TIAM1 is a RAC1-specific guanine nucleotide exchange factor (GEF) (88, 89). The majority of the cellular activity of TIAM1 is attributed to its activation of the small GTPase RAC1, switching it from an inactive GDP-bound state to an active GTP-bound state (90). TIAM1-dependent activation of RAC1 plays fundamental roles in a variety of cellular processes including cellular proliferation, migration, differentiation (91–94) and inflammation (95). In epithelial cells, TIAM1 is reported to facilitate formation of lamellipodia and filopodia (96–99). In keratinocytes, TIAM1 controls polarization of migratory cells (100–102). Loss-and gain-of-function studies have shown that TIAM1 regulates cell proliferation, survival and migration. Of relevance to our study, TIAM1 is reported to mediate neurite outgrowth, branching and angiogenesis (95, 103). Ours is the first study to examine NSC23766-mediated TIAM1-RAC1 inhibition in BPH budding and branching. Of note, NSC23766 has been reported to abrogate branching morphogenesis in other organ systems (104), and treatment with NSC23766 resulted in decreased lung branching in mouse and human explants (105, 106).

Further, previous studies on NSC23766 in the context of other pathologies have resulted in cellular toxicities at concentrations ranging from 25 µM to 800 µM. Of note, these concentrations are more than 10-fold higher (39, 107–109) compared with the concentrations we found effective in reducing the budding and branching of prostatic organoids. In addition, previous animal studies have demonstrated that NSC23766 can be applied *in vivo* without apparent evidence of organ toxicities in mice (110). In this context, our results that show reappearance of organoid budding and branching subsequent to the discontinuation of NSC23766 preclude cell death due to NSC23766 as a possible explanation for the observed reduction in budding and branching. Despite potential cytotoxicity concerns in benign cells, no significant adverse effects of NSC23766 have been reported in recent animal models (111–116).

The role of stroma in prostate pathology is well established. It has been reported previously that NSC23766 leads to a decrease in proliferation of prostatic stromal cells (117). It is hypothesized that the stroma is a crucial player in BPH budding and branching since paracrine interactions between stroma and epithelia have an established role during embryonic development of the prostate (18). However, the molecular mechanisms that drive these stromal-epithelial interactions in BPH pathogenesis remain elusive. Stromal activity plays an undeniable role in BPH pathophysiology, and the impact of stroma on therapeutic responses has been well-documented (18, 42, 43). Of note, our study shows the inhibitory effects of NSC23766 on budding and branching of prostatic organoids even in the presence of stroma.

In summary, our study using multiple approaches including bioinformatic-and experimental-based analyses points to the critical role of TIAM1 in driving the budding and branching in BPH. These results point to the idea of using TIAM1-RAC1 inhibitors as a therapeutic modality for the treatment BPH patients.

## Supporting information

Table S 3_CMap compounds

## Acknowledgments

The authors are grateful to Barry J. Maurer and Gail A. Cornwall (TTUHSC) for critical reading of the manuscript. The authors would like to thank Texas Tech University Health Sciences Center (TTUHSC) Clinical Research Institute (CRI), Lubbock, Texas for their help and support. The funding of this project was supported with startup funds from TTUHSC to MT.

**Fig. S 1:**
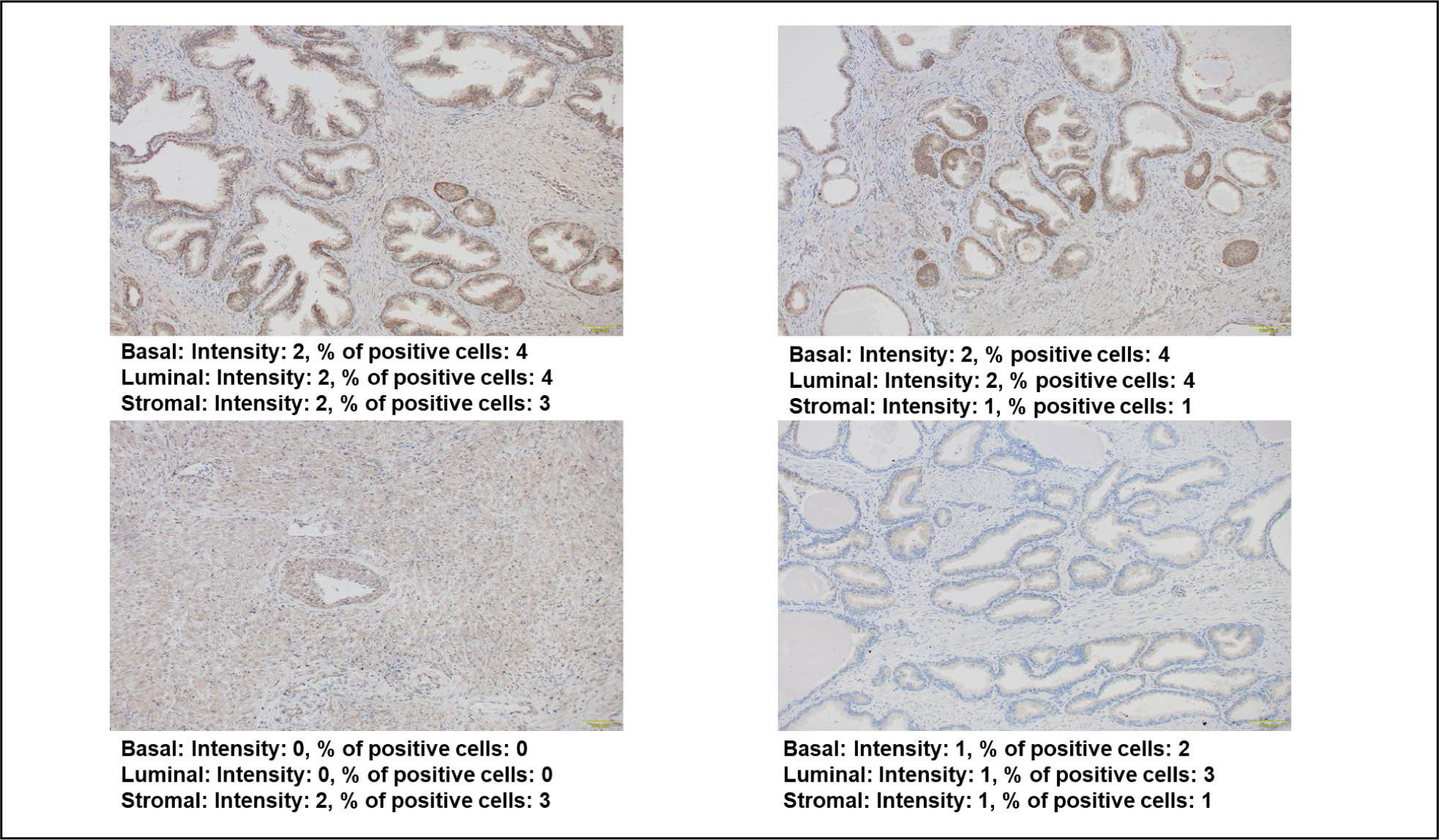
Immunohistochemical staining for TIAM1 to illustrate Immunoreactive Scoring (IRS): Representative IHC staining images of human benign prostate tissue samples with varying intensities of staining and percentage of positive cells.

**Fig. S 2.**
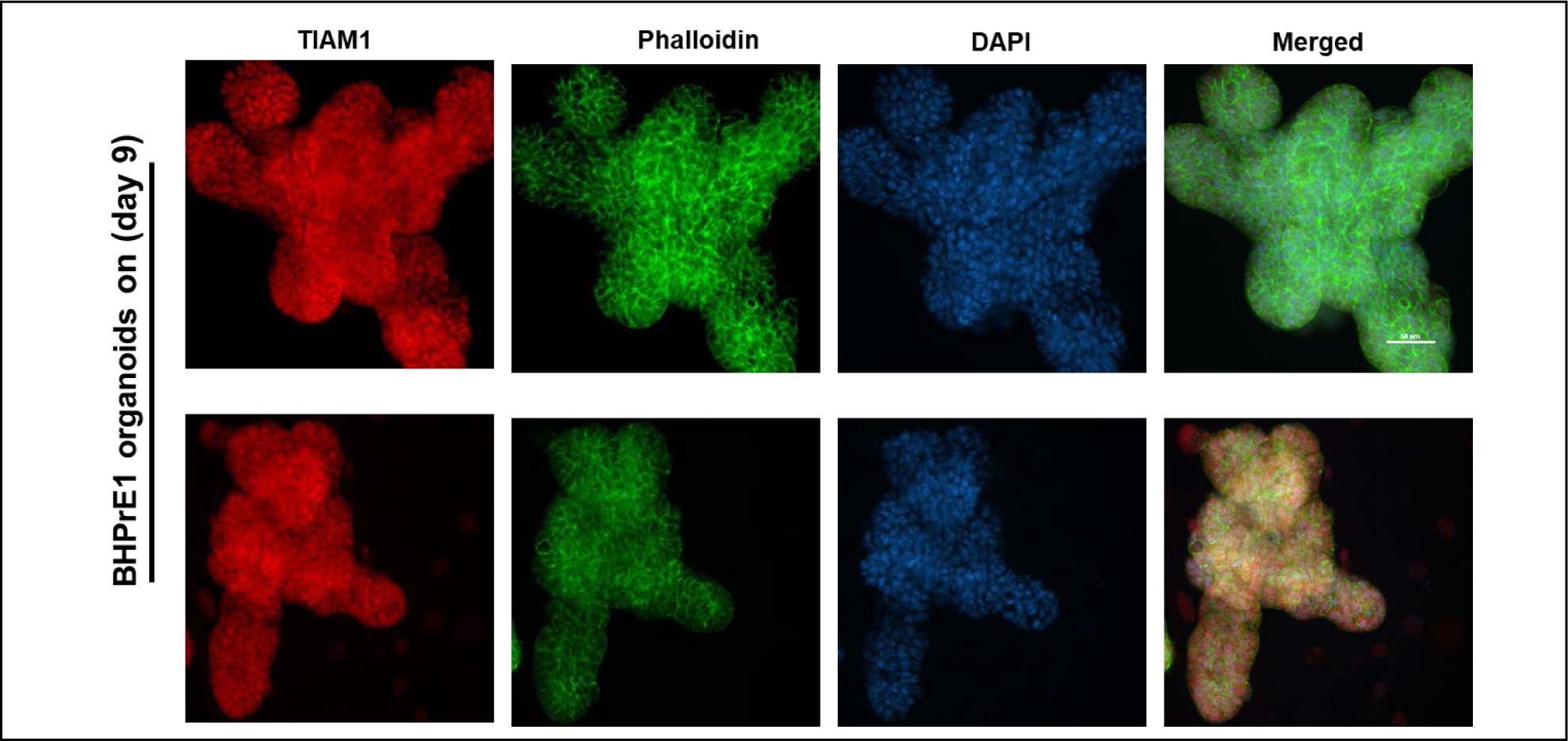
Whole-mount Immunofluorescence staining of BHPrE1 organoids stained for TIAM1 (red), Phalloidin (green) and DAPI (blue).

**Fig. S 3:**
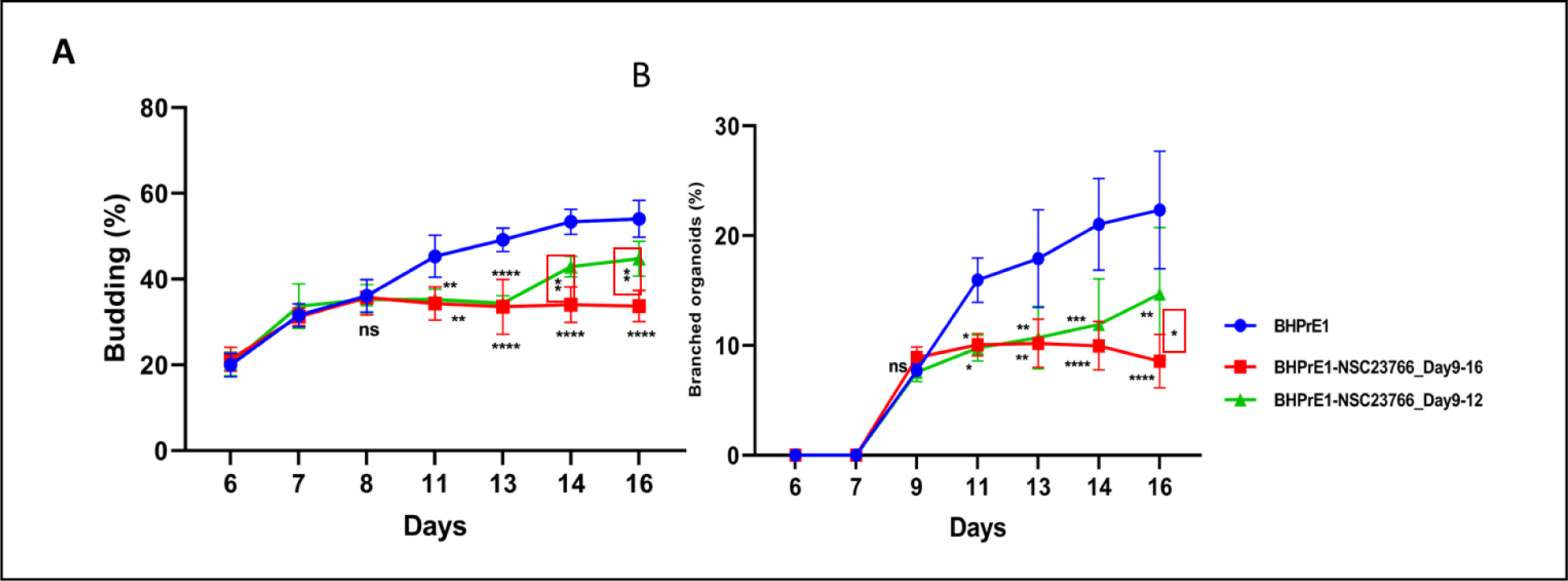
Discontinuation of NSC23766 treatment results in reinitiation of buds and branches in organoids. **A)** Quantification of the percentage of BHPrE1 budded organoids with vehicle control (blue circles), NSC23766 treatment from day 9 to day 16 (red squares) or NSC23766 treatment from day 9 to day 12 (green triangles). **B)** Quantification of the percentage of BHPrE1 branched organoids with vehicle control (blue circles), NSC23766 treatment from day 9 to day 16 (red squares) or NSC23766 treatment from day 9 to day 12 (green triangles). N = 3, Mean ± DS. Two-way ANOVA was performed, *, p < 0.05; **, p < 0.01***, p < 0.001; ****, p < 0.0001.

**Fig. S 4:**
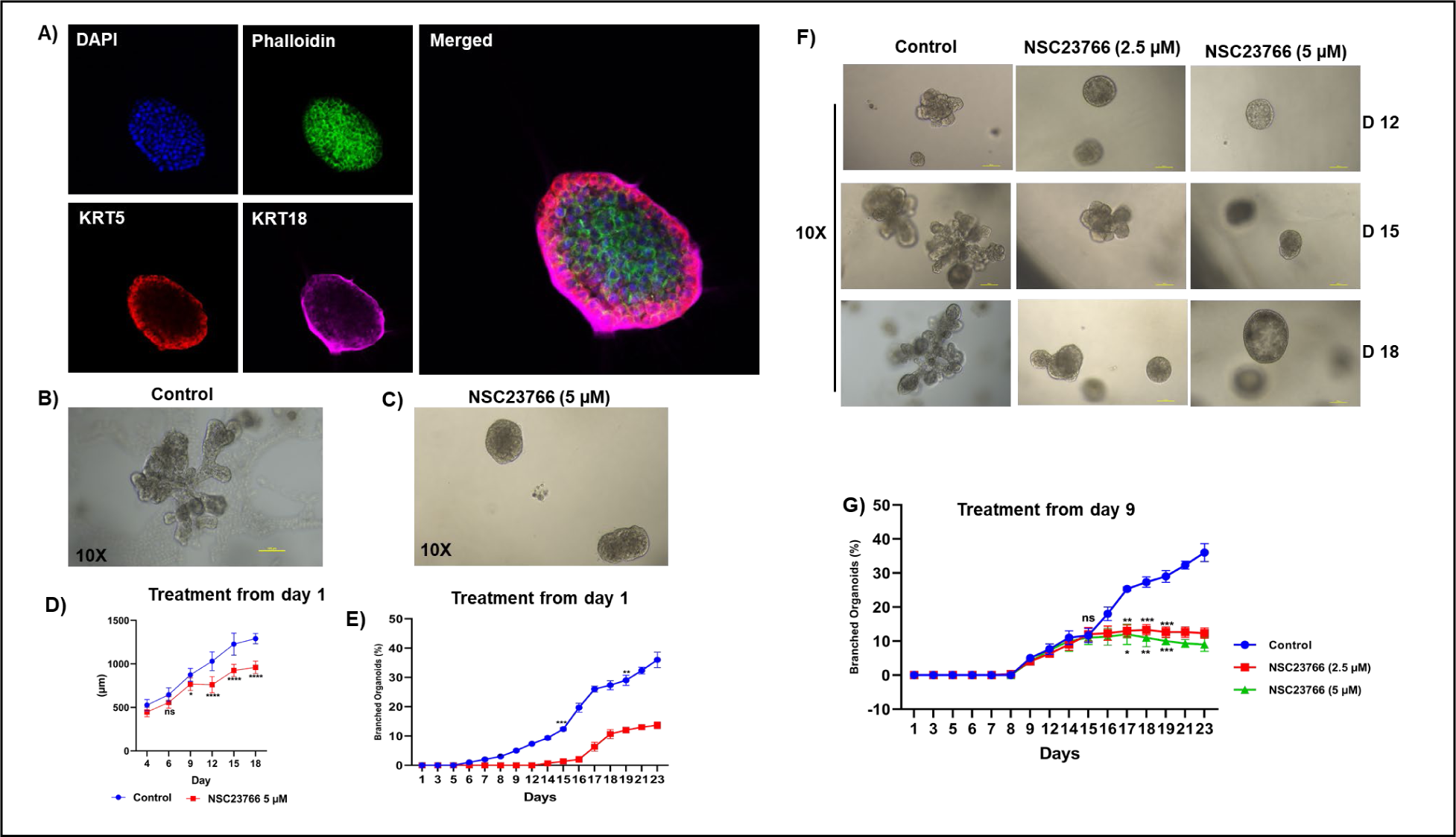
NSC23766 caused a decrease in size and branching of RWPE-1 organoids in a dose-dependent manner. **A)** Whole-mount Immunofluorescence staining of RWPE-1 organoids with basal cell marker (KRT5) and luminal cell marker (KRT8/18) on day 9. **B)** Representative images taken on day 12 of RWPE-1 organoids. Representative images taken on day 12 of RWPE-1 organoids with NSC23766 treatment starting on day 1 at 5 µM concentration. **D)** Quantification of the size of RWPE-1 organoids with NSC23766 treatment starting on day 1 at 5 µM concentrations compared with vehicle control. **E)** Quantification of the percentage of RWPE-1 branched organoids with NSC23766 treatment starting on day 1 at 5 µM concentrations compared with vehicle control. **F)** Representative images of RWPE-1 with NSC23766 treatment starting on day 1 at 2.5 µM and 5 µM concentrations compared with vehicle control. **G)** Quantification of the percentage of RWPE-1 branched organoids with NSC23766 treatment starting on day 1 at 2.5 µM and 5 µM concentrations compared with vehicle control. (N = 3, Mean ± SD, two-way ANOVA was performed, *, p < 0.05; **, p < 0.01; ***, p< 0.001; ****; p < 0.0001).

**Fig. S 5:**
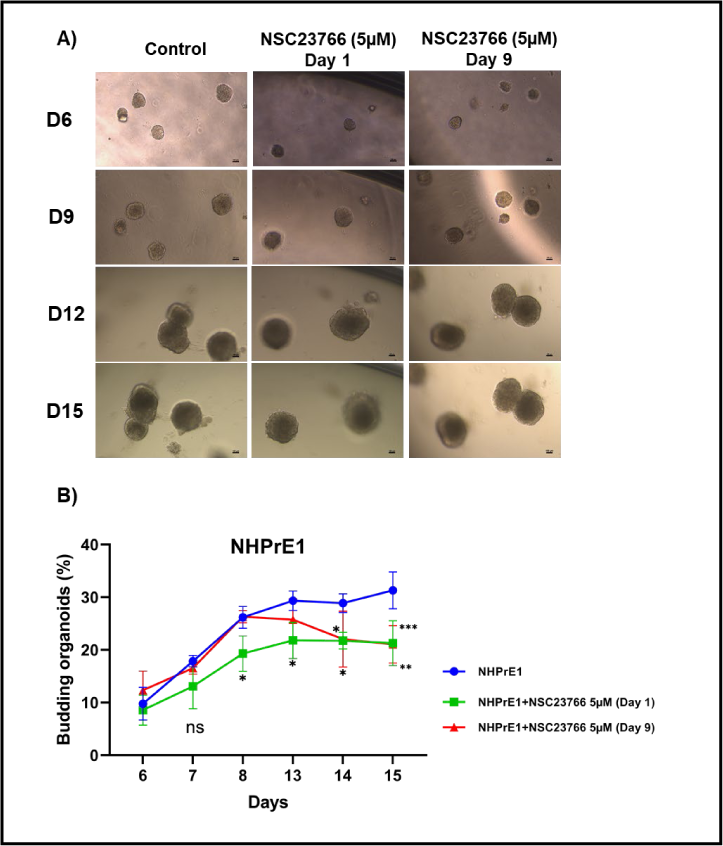
NSC23766 caused a decrease in budding of NHPrE1 organoids. **A)** Representative images taken on days 6, 9, 12 and 15 of NHPrE1 organoids treated with Vehicle control or with NSC23766 treatment on day 1 and Day 9 **B)** Quantification of the percentage of NHPrE1 budded organoids following treatment with NSC23766 on day 1 or day 9. (N=3, Mean ± SD, two-way ANOVA was performed, *, p < 0.05; **, p < 0.01; ***, p< 0.001).

**Table S 1:**
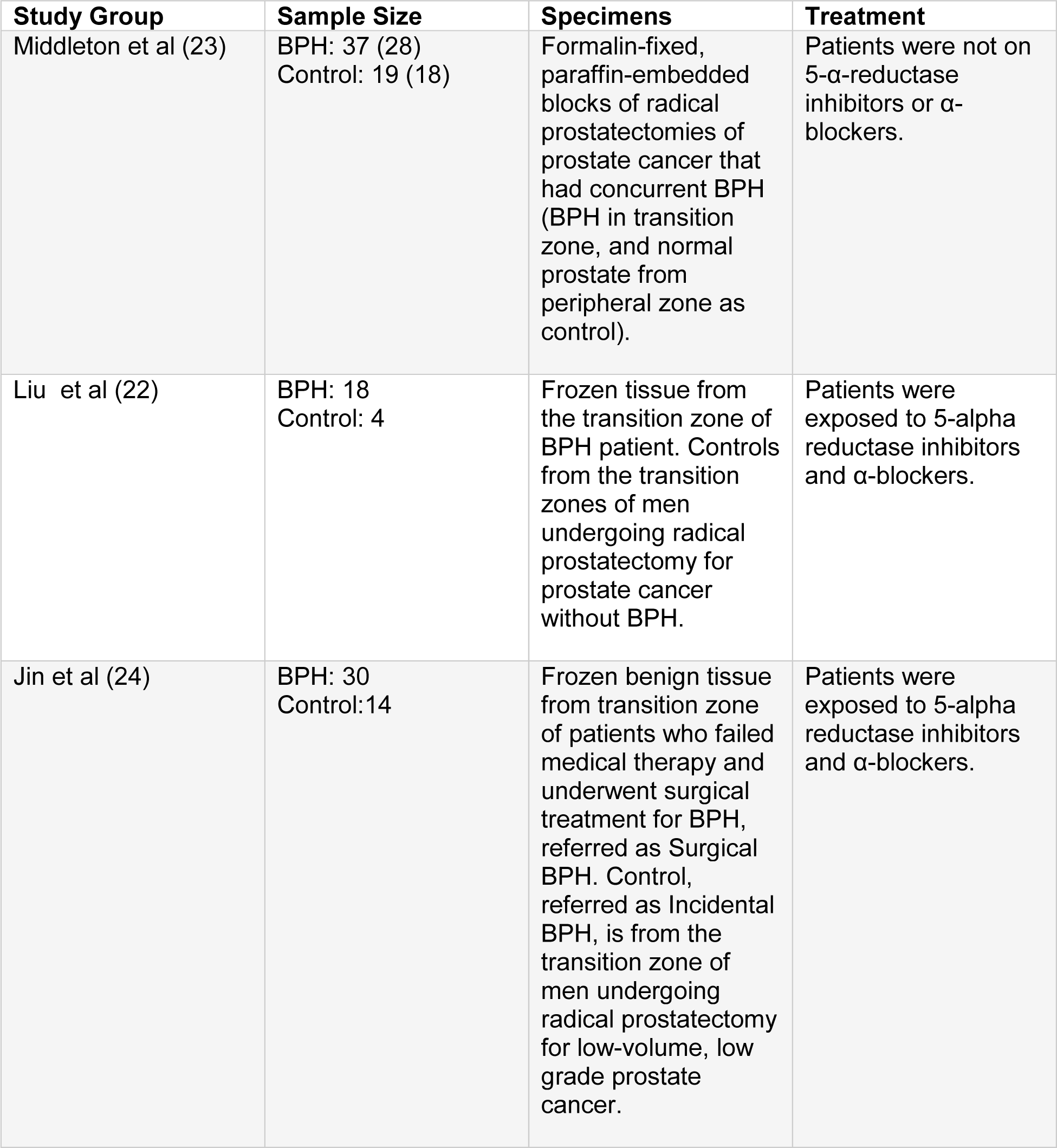
RNA-seq datasets utilized in the study for identifying the differentially expressed genes in BPH.

